# Toward reproducible, scalable, and robust data analysis across multiplex tissue imaging platforms

**DOI:** 10.1101/2020.12.11.422048

**Authors:** Erik Ames Burlingame, Jennifer Eng, Guillaume Thibault, Koei Chin, Joe W. Gray, Young Hwan Chang

**Affiliations:** Computational Biology Program, Department of Biomedical Engineering, Oregon Health and Science University, Portland, OR, USA; OHSU Center for Spatial Systems Biomedicine, Department of Biomedical Engineering, Oregon Health and Science University, Portland, OR, USA; Knight Cancer Institute, Oregon Health and Science University, Portland, OR, USA

**Keywords:** multiplex tissue imaging, GPU data science, breast cancer

## Abstract

The emergence of megascale single-cell multiplex tissue imaging (MTI) datasets necessitates reproducible, scalable, and robust tools for cell phenotyping and spatial analysis. We developed open-source, graphics processing unit (GPU)-accelerated tools for intensity normalization, phenotyping, and microenvironment characterization. We deploy the toolkit on a human breast cancer (BC) tissue microarray stained by cyclic immunofluorescence and benchmark our cell phenotypes against a published MTI dataset. Finally, we demonstrate an integrative analysis revealing BC subtype-specific features.

## 1. MAIN

Multiplex tissue imaging (MTI) methods like cyclic immunofluorescence (CyCIF) [1, 2], CODEX [3], multiplex immunohistochemistry (mIHC) [4], imaging mass cytometry (IMC) [5], and multiplex ion beam imaging [6] enable measurements of the expression and spatial distribution of tens of markers in tissues, and have facilitated our understanding of the interactions and relationships among distinct cell types in diverse tissue microenvironments. Nevertheless, for MTI to reach its full potential as a research paradigm, numerous computational challenges must be overcome, including (1) reproducible normalization of single-cell intensity measurements to enable intra- and inter-sample comparisons; (2) robust cell phenotyping at megascale to enable comparison-and soon compilation-of MTI datasets from different platforms; and (3) the development of insightful spatial features to characterize the microenvironment of the tissue or disease of interest, and so enable discrimination between tissues that vary over important clinical parameters.

To address these challenges, we present (1) a broadened application of our data-intrinsic normalization method [7], which leverages the mutually exclusive expression pattern of marker pairs in MTI stain panels to estimate normalization factors without subjective and time-consuming manual gating; (2) a distributed and graphics processing unit (GPU)-accelerated implementation of PhenoGraph [8], the popular graph-based algorithm for subpopulation detection in high-dimensional single-cell data; and (3) an integrative analysis using this toolkit on ~1.3 million cells from a 180-sample, pan-subtype human breast cancer (BC) tissue microarray (TMA) dataset (Supplementary Figure 1A-C) stained by CyCIF using a marker panel that characterizes tumor, immune, and stromal compartments (Figure 1A, Supplementary Table 1). Through consideration of both tissue composition and architecture, we identify features independent from hormone receptor (HR) and human epidermal growth factor receptor 2 (HER2) expression which discriminate between the canonical BC subtypes.

**Figure 1:**
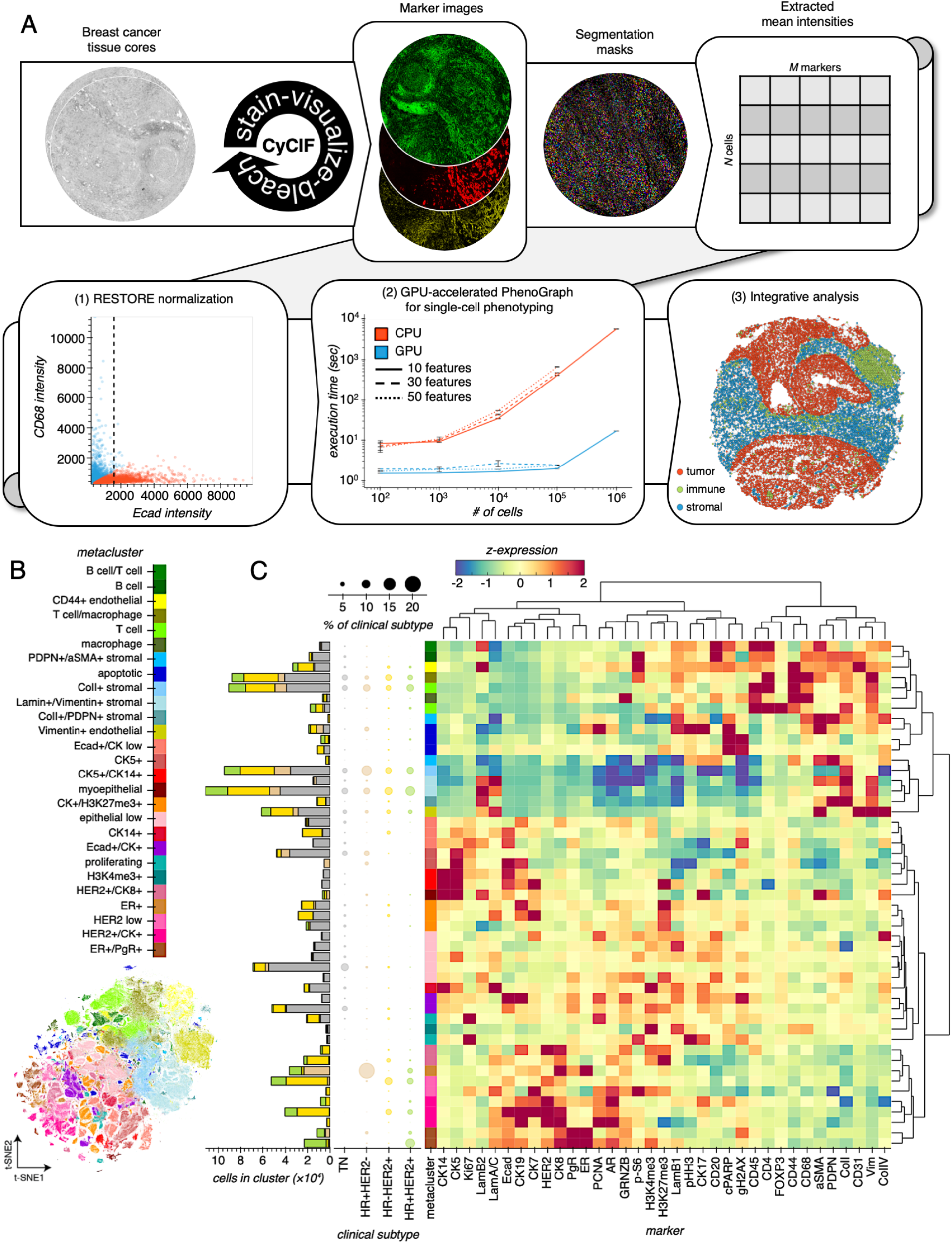
Defining single-cell phenotypes across breast cancer clinical subtypes. (A) Overview of CyCIF analysis workflow. Once TMA cores are stained by CyCIF, cells are segmented and cell mean intensities are extracted, normalized, then used to define cell phenotypes for analyses of tissue composition and architecture. Box (1) shows an example of RESTORE normalization of single-cell Ecad intensities using mutually-exclusive expression of CD68 to derive a normalization factor for Ecad. Box (2) shows benchmarking results for CPU and GPU implementations of PhenoGraph for phenotyping of simulated single-cell datasets. Compared to the legacy CPU implementation, our GPU implementation of PhenoGraph is orders of magnitude faster at scale. Error bars show standard deviation of three replicate executions. Box (3) shows the spatial layout of high-dimensional cell phenotypes in a representative tissue core. (B) t-SNE embedding of full single-cell CyCIF dataset colored by cell phenotype metacluster. (C) Hierarchical clustering of PhenoGraph clusters and CyCIF markers. The colorscale represents the z-scored marker expression. The scatter plot displays how each BC subtype is composed, where point size represents the percentage of that BC subtype that is composed of that cluster. The bar plot represents that absolute number of cells belonging to each cluster and BC subtype.

For RESTORE normalization [7] of each TMA core, we leverage the fact that tumor, immune, and stromal cells exhibit mutually exclusive expression of cell type-specific markers, and use a graph-based clustering to define positive and negative cells and normalization factors (Supplementary Figure 2A, Supplementary Table 2, see subsubsection 2.3.2). When the raw expression vectors of all cells across TMAs are embedded by t-stochastic neighbor embedding (t-SNE) [9], cells are segregated based on TMA source (Supplementary Figure 2B, left), mainly due to batch effect and in part due to subtype bias within TMAs (Supplementary Figure 1B-C). Following normalization, shared cell types between TMAs are co-embedded (Supplementary Figure 2B, right) and immune, tumor, and stromal cells are segregated (Supplementary Figure 2C), a validation of the normalization process.

To define cell types among the ~1.3 million cells in the normalized feature table, we first attempted to use the central processing unit (CPU)-based version of the widely-used algorithm PhenoGraph [8], but found it to be inefficient at this scale. To overcome this computational bottleneck, we re-implemented PhenoGraph to be executable on GPUs. Using the Python libraries RAPIDS [10] and CuPy [11] to parallelize and accelerate several of PhenoGraph’s computations (see subsection 2.4), we observed multiple orders of magnitude improvement in the algorithm’s speed without sacrificing clustering quality (Figure 1A, box (2)). Our PhenoGraph implementation identified diverse tumor, immune, and stromal cell types across tissues and BC subtypes (Figure 1B-C). To define phenotypes shared across tissues, metaclusters of similar phenotypes were aggregated based on the hierarchical clustering of phenotypes based on their mean marker expression. While tissues from all BC subtypes contained similar populations of immune, stromal, and endothelial cells, differences between BC subtypes were largely driven by variable tumor cells expression of luminal and basal cytokeratins, HER2, and the hormone receptors ER and PgR, as previously reported [12]. We further assessed the robustness of our identified cell phenotypes by either subsampling tissue cores (Supplementary Figure 3A-C), or by applying ±20% noise to normalization factors for each marker for each core (Supplementary Figure 3D) and found that phenotypes are identified as they are sampled and are robust to minor variation in normalization factors.

With the growth of MTI in the cancer research and translational communities, there is an acute need for robust and integrative analyses of MTI data across platforms and cohorts [13]. In a step toward addressing that need, we validated our identified cell types through comparison with a recently published survey of BC by IMC [12]. While the total number of cells from each BC subtype varied between the Basel (IMC) and OHSU (CyCIF) cohorts (Supplementary Figure 4A), there was substantial overlap between the marker panels used for each MTI platform (Supplementary Figure 4B). By aligning the cell phenotypes independently detected by PhenoGraph in each cohort (Supplementary Figure 4C), we found highly-correlated clusters for stromal, immune, basal, and proliferating cell types, among others (Supplementary Figure 4D), suggesting that shared cell types could be matched across cohorts and MTI platforms, a necessary step for data integration. We note that differences between cohort cell types may reflect the differences between cohort composition with respect to BC subtype. Consistent between cohorts and platforms, tumor cells differed more between samples than did immune, stromal, and endothelial cells (Supplementary Figure 5).

Although BC is appreciated as a genetically and morphologically heterogeneous disease, its clinical subtyping is based on the expression of relatively few markers—in particular, tumor cell expression of the hormone receptors for estrogen and progesterone (ER and PgR, respectively, or HR, collectively), and human epidermal growth factor 2 (HER2)—which is insufficient to explain differences in treatment response within each subtype [14]. Recent studies using MTI to interrogate intact BC tissues have found that the spatial contexture of the BC microenvironment can improve our ability to predict clinical outcome [12, 15]. However, because these studies have either focused on disease risk or a single BC clinical subtype, here we focused on compositional and spatial features which differentiate between subtypes.

At the composition level, we first considered the tumor cell differentiation states of BC subtypes through their expression of luminal and basal cytokeratins (CKs). While CK+ cells in HR-/HER2+, HR+/HER2-, and HR+/HER2+ tissues primarily expressed luminal CKs 19, 8, and 7, CK+ cells in TN tissues exhibited significantly greater differentiation state heterogeneity (Figure 2A), as they expressed many different combinations of luminal and basal cytokeratins (Figure 2B). This differentiation state heterogeneity is consistent with the genetic and histological heterogeneity of triple negative BC (TNBC) described in other studies [16, 17]. We next determined the composition of each tissue core with respect to the cell metaclusters we defined above. Hierarchical clustering of cores based on their cell metacluster densities highlighted the broad variability of cellular composition within and between BC subtypes (Figure 2C-D). When the cell metaclusters were further aggregated into immune, stromal, and tumor cell types (see subsubsection 2.8.2), we found the HR+HER2-tissues to have lower overall immune cell density than the other BC subtypes, and no differences in stromal or tumor cell density between subtypes (Figure 2E).

**Figure 2:**
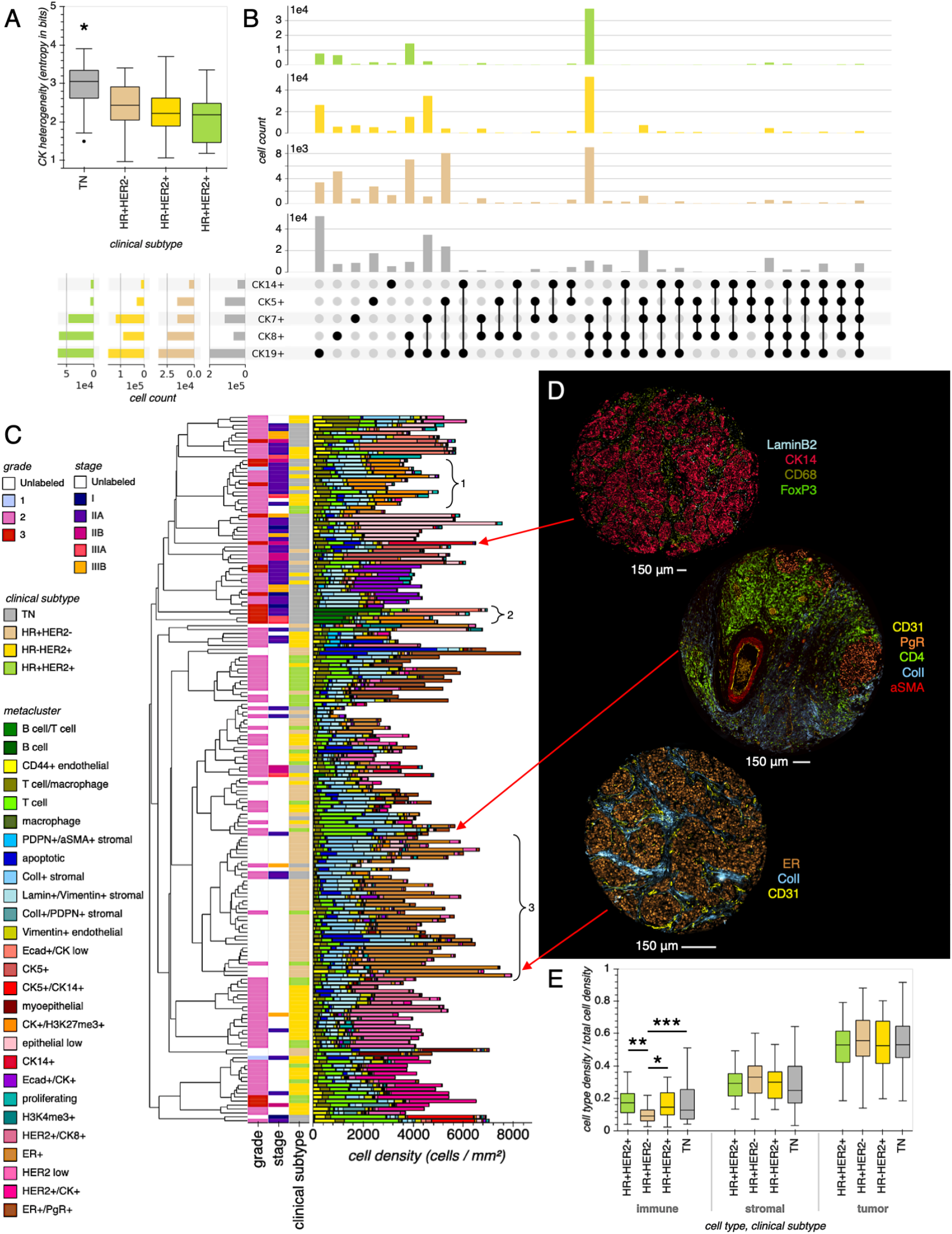
Breast cancer subtypes are differentiated by single-cell composition. (A) Epithelial differentiation heterogeneity across BC subtypes. Box plot displaying cytokeratin (CK) expression heterogeneity across BC subtypes, where each box represents the distribution of tissue cores from a BC subtype, and each core is summarized based on the entropy of the distribution of CK+ cell types contained within it. Group-wise comparisons were made using one-way ANOVA with pairwise Tukey post-hoc test (TN, *n* = 47; HR+HER2-, *n* = 52; HR-HER2+, n = 53; HR+HER2+, n = 28). *P < 0.001 for all TN comparisons with other BC subtypes. (B) UpSet plot summarizing the distribution of CK+ cell types across BC subtypes, considering each CK alone (left margin) or in combination (upper margin). (C) Cell phenotype density across tissue cores. Bar plot where each bar represents a TMA core, the full bar height represents its total cell density, and each colored segment represents the density of a particular cell metacluster. Bars are hierarchically clustered based on cell metacluster densities. Each bar is labeled with its corresponding subtype, stage, and grade, if a label is available. The inset brackets indicate (1) cores with abundant H3K27me3+ tumor cells, which could indicate a mechanism of HR repression in some TN and HR-/HER2+ tissues [31]; (2) cores with abundant infiltrating B cells (TIL-B), consistent with association found between TIL-B and high-grade, HR-BC [32]; and (3) cores with relatively low immune density, consistent with the finding that HR+/HER-tissues are immunologically cold compared to TN and HER2+ tissues [33, 34]. (D) A selection of representative tissue cores. (E) The immune, stromal, and tumor densities of tissue cores from each BC subtype. Groupwise comparisons were made using one-way Welch ANOVA and Games-Howell post-hoc test. *P = 0.034, **P = 0.035, ***P = 0.079 (TN, n = 47; HR+HER2-, n = 52; HR-HER2+, n = 53; HR+HER2+, n = 28).

Recognizing that cell density measurements fail to capture the organization of cells in each tissue, we next characterized the spatial architectures of BC subtypes by building cell neighborhood graphs for each tissue. Given the recent evidence that the quantity and diversity of BC tumor cell interactions with other cell types can inform disease outcome [12, 15], we first identified the neighboring cells to each tumor cell and compared the composition of tumor cell neighborhoods across BC subtypes (Figure 3A). Though most tumor cell interactions (~70-80%) are with other tumor cells typical to their BC subtype, we observed increased tumor-stromal interaction in the HR+HER2-subtype. When considering tissues that contain an appreciable population of stromal cells (tissues comprised of at least 25% stromal cells), we confirmed that there was significantly more stromal mixing with tumor cells in HR+HER2-tissues (Figure 3B). Importantly, stromal mixing can vary widely between tissues in spite of their similar stromal density (Figure 3C), highlighting the importance of the spatial context that is preserved in intact tissues. Since malignant epithelial cells can suppress fibroblast maturation and thus promote fibroblast aromatase activity [18], ER+HER2-tumors likely favor more from proximal fibroblasts as a source of growth-inducing estrogen than other BC subtypes, and may even act to maintain tumor microenvironments with high stromal mixing [19].

**Figure 3:**
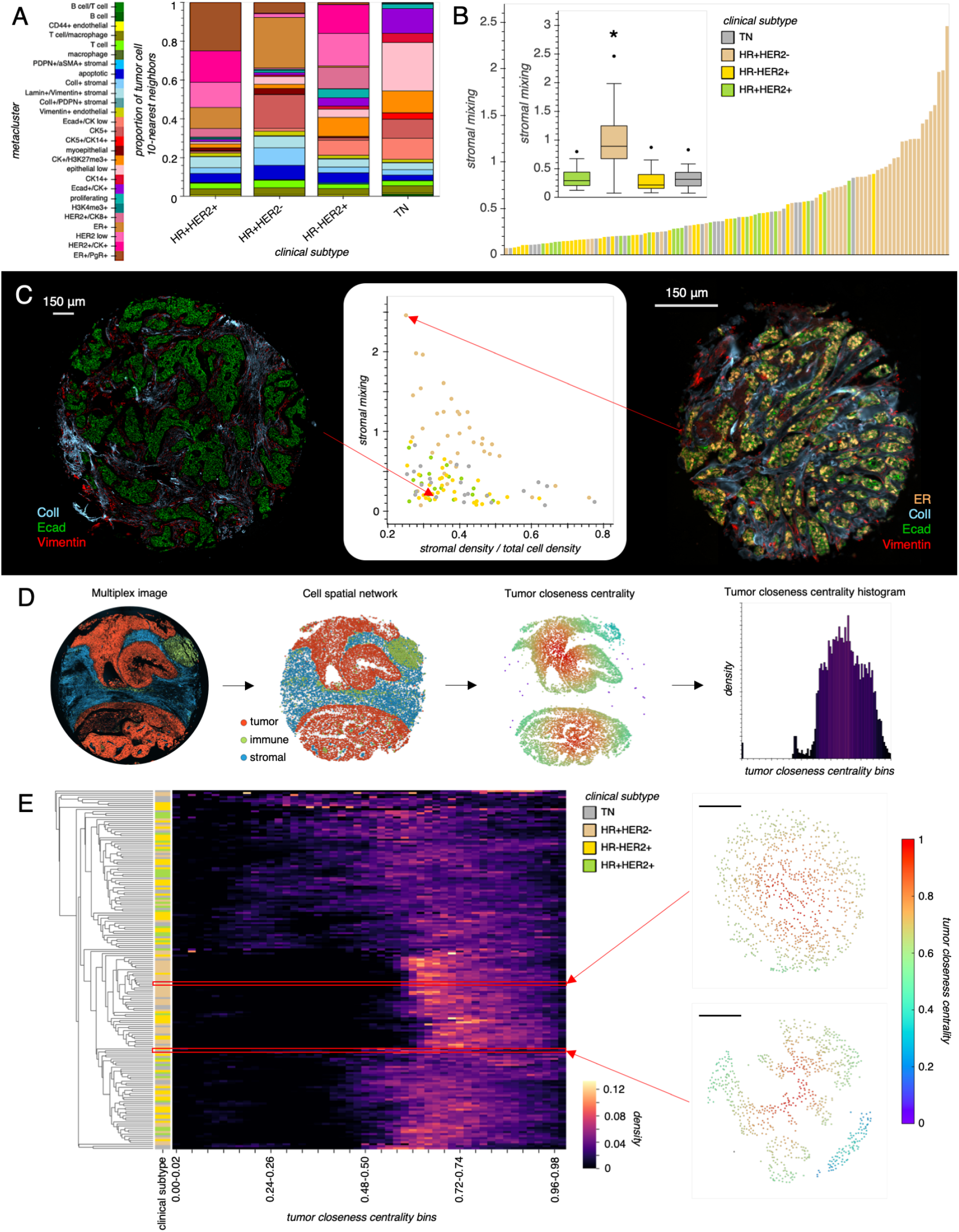
Breast cancer cellular composition belies tumor-stromal interaction and tumor architecture. (A) Stacked bar plots displaying the proportion of tumor cell 10-nearest neighbors for each BC subtype. Each colored bar segment represents the proportion of tumor cell neighbors that are comprised of the corresponding cell metacluster. (B) Bar plot representation of tissue core stromal mixing, where only cores with greater than 0.25 stromal fraction are shown. Cores are oredered based on increasing stromal mixing. Inset shows box plot comparing stromal mixing over BC subtype. Groupwise comparisons were made using one-way Welch ANOVA and Games-Howell post-hoc test. *\*P* <0.001 for all HR+HER2-comparisons (TN, n = 47; HR+HER2-, n = 52; HR-HER2+, n = 53; HR+HER2+, n = 28). (C) Scatter plot displaying stromal density versus stromal mixing and images of representative cores with similar stromal density but different stromal mixing. (D) Overview of tumor architecture characterization. A spatial graph is defined over tumor cells, where each tumor cell is connected to others within a 65 μm radius from its centroid, and the closeness centrality is then measured over this graph. Each core is then summarized as a histogram of centrality values. (E) Hierarchical clustering of cores based on their tumor closeness centrality histograms. The upper and lower outset tumor graphs correspond to the right and left tissue core images in Figure 3C, respectively. Scale bar in tumor graphs is 150 μm.

We reasoned that differences in tumor-stromal interaction might translate into detectable differences between BC subtypes based on their tumor architectures alone. To characterize the tumor architecture of each tissue, we constructed tumor architecture graphs over which we computed the closeness centrality for each tumor cell, which quantifies the relative closeness of that tumor cell to all other tumor cells in the tissue (Figure 3D). Consistent with the stromal mixing trend observed above, tumor cells in HR+HER2-tissues tended to have increased closeness centrality compared to tumor cells from the other BC subtypes (Figure 3E), which is in part a reflection of HR+HER2-tumor cell nests tending to be separated by narrower streams of stromal cells than tumor nests in tissues from other BC subtypes (Figure 3C). In summary, by analyzing BC tissues with spatially resolved MTI, we have identified inter- and intra-cell phenotype interactions which can be leveraged to help discriminate between canonical BC subtypes on a basis other than receptor expression.

This work is motivated by an understanding that the spatial context of the tumor microenvironment in intact cancer tissues enables a more granular definition of disease, and—we hope—the design of more personalized and effective treatments. With spatially-resolved MTI, our analysis makes clear that the cellular composition of BC tissue can belie important aspects of its spatial architecture. Ongoing work involves validating these findings in a cohort with more extensive clinical annotation to assess their significance to disease outcome between and within BC subtypes. While BC cell phenotypes and architectural features we have derived will be assets to future BC studies, our generic toolkit can be used standalone or integrated with existing toolkits [20] to improve the efficiency and reproducibility of analytics for any single-cell measurement platform.

## 2. METHODS

### 2.1. Acquisition of breast cancer tissue microarrays (TMAs)

The tissues used in this study are a compilation of multiple TMAs: BR1201a-SG48 (US Biomax Inc., https://www.biomax.us/BR1201at), BR1506-A019 (US Biomax Inc., https://www.biomax.us/tissue-arrays/Breast/BR1506), Her2B-K154 (US Biomax Inc., https://www.biomax.us/tissue-arrays/Breast/Her2B), and the TransATAC TMAs T-ATAC-4A-Left and T-ATAC-4A-Right [21]. All tissues that were successfully stained and imaged were included in the study, representing 180 tissue cores from 128 patients.

### 2.2. Cyclic immunofluorescence (CyCIF) staining of tissues

#### 2.2.1. Tissue preparation

Formalin-fixed paraffin-embedded (FFPE) human tissues were received mounted on adhesive slides. The slides were baked overnight in an oven at 55 °C (Robbin Scientific, Model 1000) and an additional 30 minutes at 65 °C (Clinical Scientific Equipment, NO. 100). Tissues were deparaffinized with xylene and rehydrated with graded ethanol baths. Two step antigen retrieval was performed in the Decloaking Chamber (Biocare Medical) using the following settings: set point 1 (SP1), 125 °C, 30 seconds; SP2: 90 °C, 30 seconds; SP limit: 10 °C. Slides were further incubated in hot Target Retrieval Solution, pH 9 (Agilent, S236784-2) for 15 minutes. Slides were then washed in two brief changes of diH2O (~2 seconds) and once for 5 minutes in 1x phosphate buffered saline (PBS), pH 7.4 (Fisher, BP39920). Sections were blocked in 10% normal goat serum (NGS, Vector S-1000), 1% bovine serum albumin (BSA, Sigma A7906) in PBS for 30 minutes at 20 ^°^C in a humid chamber, followed by PBS washes. Primary antibodies were diluted in 5% NGS, 1% BSA in 1x PBS and applied overnight at 4 ^°^C in a humid chamber, covered with plastic coverslips (Bio-Rad, SLF0601). Following overnight incubation, tissues were washed 3 x 10 min in 1x PBS. Coverslips (Corning; 2980-243 or 2980-245) were mounted in Slowfade Gold plus DAPI mounting media (Life Technologies, S36938).

#### 2.2.2. Fluorescence microscopy

Fluorescently stained slides were scanned on the Zeiss AxioScan.Z1 (Zeiss, Germany) with a Colibri 7 light source (Zeiss). The filter cubes used for image collection were DAPI (Zeiss 96 HE), Alexa Fluor 488 (AF488, Zeiss 38 HE), AF555 (Zeiss 43 HE), AF647 (Zeiss 50) and AF750 (Chroma 49007 ET Cy7). The exposure time was determined individually for each slide and stain to ensure good dynamic range but not saturation. Full tissue scans were taken with the 20x objective (Plan-Apochromat 0.8NA WD=0.55, Zeiss) and stitching was performed in Zen Blue image acquisition software (Zeiss).

#### 2.2.3. Quenching fluorescence signal

After successful scanning, slides were soaked in 1x PBS for 10-30 minutes in a glass Coplin jar, waiting until glass coverslip slid off without agitation. Quenching solution containing 20 mM sodium hydroxide (NaOH) and 3% hydrogen peroxide (H2O2) in 1x PBS was freshly prepared from stock solutions of 5 M NaOH and 30% H2O2, and each slide placed in 10 ml quenching solution. Slides were quenched under incandescent light, for 30 minutes for FFPE tissue slides. Slides were then removed from chamber with forceps and washed 3 x 2 min in 1x PBS. The next round of primary antibodies was applied, diluted in blocking buffer as previously described, and imaging and quenching were repeated over ten rounds for FFPE tissue slides.

### 2.3. Data pre-processing

#### 2.3.1. Cell segmentation and mean intensity extraction

Cell segmentation and mean intensity extraction were performed as previously described [2]. Mean intensities for each cell were extracted from the biologically-relevant compartment for each marker, i.e. mean intensities for markers with known nuclear (cytoplasmic) localization were extracted from nuclear (cytoplasmic) segmentation masks (Supplementary Table 2). Cytoplasmic segmentation masks were computed by subtracting nuclear segmentation masks from full cell body segmentation masks.

#### 2.3.2. Single-cell intensity normalization

Normalization factors for single-cell mean intensities were computed as previously described [7] using the putative mutually-exclusive marker pairs in Supplementary Table 2. Normalization factors are computed for each pair of reference and mutually-exclusive markers, and the median of these factors is used to normalize each raw single-cell mean intensity vector for each CyCIF marker and each TMA core. Raw intensities were normalized using the equation:

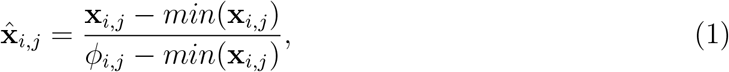

where 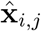, and **x**_*i,j*_ are the normalized and raw single-cell mean intensity vectors for CyCIF marker *i* for all cells in tissue core *j*, respectively, and *ϕ_i,j_* is the corresponding normalization factor determined as described above. Therefore, cells with a normalized intensity greater than 1 are considered to be above the background intensity level.

### 2.4. Single-cell phenotyping

#### 2.4.1. GPU acceleration of PhenoGraph

Given a cell-by-feature dataframe, the PhenoGraph algorithm [8] consists of two primary steps: (1) defining a k-nearest neighbor graph over all cells that is then refined by computing the Jaccard similarity measure over graph edges, and (2) partitioning the graph into discrete cell phenotypes through optimization of partition modularity, such that cells in the same partition are more connected to each other than to cells of another partition. In the official version of PhenoGraph (https://github.com/dpeerlab/PhenoGraph), these steps are implemented using a combination of Python and C++ libraries that execute on CPU. In Figure 1A, we show that PhenoGraph execution time increases exponentially with increasing dataset size, taking approximately 3 hours to process a synthetic 1 million cell-by-10 feature dataset. Most MTI datasets measure tens of features, but CPU-based PhenoGraph was unable to fully process the 1 million cell-by-30- and 50-feature synthetic datasets in the 8 hours allotted for the experiment. We see such computational bottlenecks-which would be even further constricted when compiling multiple MTI or cytometry datasets-as a major obstacle to current studies and future meta-studies of high-dimensional MTI datasets, where rapid iteration will be essential to the validation of cross-platform data integration techniques.

Owing to recent advances in GPU computing and its ever-broadening adoption in machine learning research, there now exist accelerated GPU-based analogs of many Python scientific computing libraries [10, 11], including those with which the CPU-based PhenoGraph is implemented. Some of these libraries even allow computation to be distributed across multiple GPUs [10, 22]. We employed two of such libraries, CuPy [11] and RAPIDS [10], to accelerate each step of the PhenoGraph algorithm and enable distributed computing over multiple GPUs. With our GPU-based implementation, it is now possible to phenotype cells in megascale cytometry datasets in seconds-to-minutes rather than hours-to-days, and without subsampling. Installation instructions and additional details about our implementation can be found at https://gitlab.com/eburling/grapheno.

#### 2.4.2. Phenotyping and metacluster annotation

Apart from the benchmarking experiment described in Figure 1A, we use only our GPU-based implementation of PhenoGraph throughout this work. Single-cell phenotypes were defined based on single-cell mean intensity for the 35-marker CyCIF panel (Supplementary Table 1). Prior to application of PhenoGraph, data were 99.9th-percentile normalized and arcsin transformed (cofactor = 5). Following [12], PhenoGraph was parameterized (k=40) to over-cluster the data and detect rare cell types. PhenoGraph clustering was followed by aggregation of phenotypes into metaclusters based on hierarchical clustering of phenotype mean marker intensities and to preserve known biological variation.

#### 2.4.3. Robustness of derived cell phenotypes

To assess PhenoGraph clustering robustness to sampling shift, PhenoGraph clusters were derived using random subsets of tissues of varying cardinality, from 10% to 90% of all tissues, and compared the z-scored mean marker intensities of the PhenoGraph-derived clusters from the full reference dataset and each subset using pairwise Pearson’s correlation. Even with heavy subsampling, the median of the maximum correlations between matching clusters from reference-to-sample comparisons held at ~0.75 (Supplementary Figure 3A), indicating that we are defining a robust core set of cell phenotypes. Indeed, the major variation between reference and subsample clusters appeared to be sample-specific tumor cell phenotypes from the tissues held out from each subsample (Supplementary Figure 3B-C), suggesting that PhenoGraph defines robust cell phenotypes that are shared across tissues and is capable of detecting new phenotypes as they are added to the dataset. To assess the robustness of our derived phenotypes to variability in normalization, we also simulated ±20% measurement noise by multiplying the normalized cell intensity vectors for each tissue and marker by a scaling factor drawn uniformly at random from the range [0.8,1.2] and compared the z-scored mean marker intensities of the PhenoGraph-derived clusters from the clean reference and noisy datasets using pairwise Pearson’s correlation. Even with these significant perturbations to the intensity profiles of cells, the median of the maximum correlations between matching clean and noisy clusters held above 0.8 (Supplementary Figure 3D), indicating cluster robustness to differentials in preanalytical variables like tissue fixation or autofluorescence which can affect measured IF intensity across a TMA.

### 2.5. t-stochastic neighbor embedding (t-SNE)

To enable visualization, the full 35-feature single-cell dataset was reduced to 2 dimensions using the RAPIDS implementation of *t*-SNE [9] with default parameters except perplexity = 60. Prior to t-SNE processing, data were 99.9th-percentile normalized and arcsin transformed (cofactor = 5). Plots containing t-SNE embeddings of the full ~1.3 million-cell dataset were created using Datashader (https://github.com/holoviz/datashader).

### 2.6. Cross-platform breast cancer cell phenotype validation

The imaging mass cytometry (IMC) dataset [12] used for our cell phenotype validation experiment was retrieved from https://zenodo.org/record/3518284. To make a fair comparison between the OHSU (CyCIF) and Basel (IMC) datasets, we independently ran PhenoGraph on each using the same parameters (*k*=40, Louvain partitioning) and only the overlapping features between IMC and CyCIF stain panels (Supplementary Figure 4B). The phenotypes derived from each platform were then cross-correlated to identify inter-platform phenotype matches. Using the clustermap function from seaborn [23], the cross-correlation matrix was then hierarchically clustered with Ward linkage and used to sort the matching clusters between the two cohort heatmaps.

### 2.7. Statistical analyses

For the groupwise comparisons in Figure 2A, Figure 2E, Figure 3B, we first tested the assumption of homogeneity of variances using the bartlett function from the Python package SciPy [24]. When the assumption was (not) met, we made groupwise comparisons using one-way ANOVA with Tukey-HSD post-hoc test using the pairwise_tukey (one-way Welch ANOVA with Games-Howell post-hoc test using the pairwise_gameshowell) function from the Python package pingouin [25].

### 2.8. Tissue composition analyses

#### 2.8.1. Epithelial differentiation heterogeneity

Cells from each core were labeled as positive for each cytokeratin if their mean intensity was greater than the normalization factor computed for that cytokeratin for that core. The plot from Figure 2B was generated using UpSetPlot (https://github.com/jnothman/UpSetPlot). The CK heterogeneity of each core was computed by measuring the Shannon entropy of its distribution of CK-expressing cells. The homogeneity of variances assumption was met, so comparison of the CK heterogeneity over BC subtypes was made using the pairwise_tukey function from pingouin [25].

#### 2.8.2. Aggregation of immune, stromal, and tumor cell phenotypes

To enable high-level comparison of cell phenotype distribution over BC subtypes (Figure 2E and Figure 3), cell metaclusters were aggregated into immune, stromal, and tumor groups with the following metacluster membership:

- immune = [B cell/T cell, B cell, T cell/macrophage, T cell, macrophage]
- stromal = [CD44+ endothelial, PDPN+/aSMA+ stromal, ColI+ stromal, Lamin+/Vimentin+ stromal, ColI+/PDPN+ stromal, Vimentin+ stromal, Vimentin+ endothelial]
- tumor = [Ecad+/CK low, CK5+, CK5+/CK14+, myoepithelial, CK+/H3K27me3+, epithelial low, CK14+, Ecad+/CK+, proliferating, apoptotic, H3K4me3+, HER2+/CK8+, ER+, HER2 low, HER2+/CK+, ER+/PgR+]

#### 2.8.3. Cell phenotype density

The density of each of the 27 cell metaclusters in each tissue core was measured by counting the number of cells of each metacluster in the core, then dividing the count by the area of the convex hull defined by the cell centroids of the core. Tissue cores were then hierarchically clustered based on their z-scored cell metacluster densities using the clustermap function with Ward linkage from seaborn [23]. The homogeneity of variances assumption was not met, so comparisons of immune, stromal, and tumor cell densities over BC subtypes were made using the pairwise_gameshowell function from pingouin [25].

### 2.9. Tissue architecture analyses

#### 2.9.1. Tumor cell neighborhood interactions

To characterize the microenvironments of tumor cells across BC subtypes, we identified the cell metaclusters of the 10 nearest cells within 65 μm (double the median of the minimum tumor-stromal distances across all tissue cores) of each tumor cell. Tumor cells were then split based on the BC subtype of the tissue from which they were derived, and counts for each metacluster were summed over all tumor cells such that each metacluster could be represented as a proportion of the total tumor neighborhood for each BC subtype.

To measure the extent of tumor-stromal cell interactions in each tissue core, we computed their stromal mixing scores, an adaptation of a previously described cell-cell mixing score [15]. To focus on cores that had substantial stromal composition, we first selected cores which are comprised of at least 25% stromal cells and for each we defined a 10-nearest neighbor spatial graph over all cells in that core. Second, we removed edges between cells with an interaction distance greater than 65 μm. Finally, we computed the stromal mixing score for tissue core *j* as:

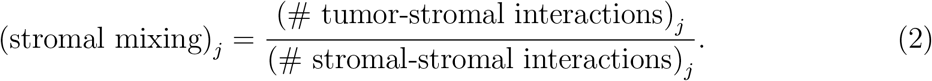

#### 2.9.2. Tumor graph centrality

To characterize tumor architecture in each tissue core, we considered the spatial interactions between tumor cells only. To account for variation in tissue core diameter (Supplementary Figure 1A) which would affect the scale of spatial graph characteristics, we subsampled large diameter cores to be equal in size to the smallest diameter cores by only considering cells within the 300 μm-radius circle drawn about the centroid of each core. With the spatially-subsampled cores, we first construct a 4-nearest neighbor spatial graph over all tumor cells in each core. Here we use *k* = 4 rather than *k* = 10 to construct a sparser graph since we are focusing on tumor cells only. Over this graph we compute the Wasserman-Faust closeness centrality of each cell using the closeness_centrality function from the Python package networkx [26]. The Wasserman-Faust closeness centrality of cell *u* is computed as:

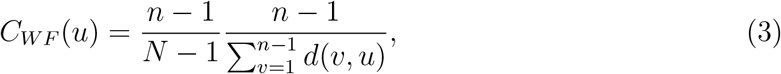

where *d*(*v,u*) is the shortest-path distance between cells *v* and *u, n* is the number of cells that can reach *u*, and *N* is the number of cells in the graph. For heatmap visualization, the distribution of tumor cell centrality for each core was max-normalized, converted into a 50-bin histogram over range = (0,1), then hierarchically clustered using the clustermap function from seaborn [23] with Jensen-Shannon distance and average linkage.

### 2.10. Plotting and visualization

Unless otherwise noted, all plots were generated using Holoviews [27] with either the Bokeh [28] or matplotlib [29] backends. Images of CyCIF-stained tissue cores were generated using napari [30].

## 2.11. Data availability

Data and code with be made available through zenodo.com and gitlab.com upon publication.

## 3. Acknowledgements

This work was supported in part by the National Cancer Institute (U54CA209988, U2CCA233280, U01 CA224012), the OHSU Center for Spatial Systems Biomedicine, and a Biomedical Innovation Program Award from the Oregon Clinical and Translational Research Institute. We acknowledge expert technical assistance by staff in the Advanced Multiscale Microscopy Shared Resource, supported by the OHSU Knight Cancer Institute (NIH P30 CA069533) and the Office of the Senior Vice President for Research. Equipment purchases included support by the OHSU Center for Spatial Systems Biomedicine, the MJ Murdock Charitable Trust, and the Collins Foundation. The resources of the Exacloud high performance computing environment developed jointly by OHSU and Intel and the technical support of the OHSU Advanced Computing Center are gratefully acknowledged. E.A.B. receives support from a scholar award provided by the ARCS Foundation Oregon.

## 4. Contributions

Conceptualization, E.A.B., J.E., G.T., K.C., J.W.G. and Y.H.C.; Methodology, E.A.B, J.E., G.T., K.C., J.W.G. and Y.H.C.; Software, E.A.B., G.T., J.E. and Y.H.C.; Validation, E.A.B., J.E., K.C., and Y.H.C.; Formal Analysis, E.A.B.; Investigation, E.A.B. and J.E.; Resources, K.C. and J.W.G.; Data Curation, E.A.B., J.E., and K.C.; Writing-Original Draft, E.A.B. and Y.H.C.; Writing-Review and Editing, E.A.B., K.C., J.W.G. and Y.H.C.; Visualization, E.A.B; Supervision, K.C., J.W.G, and Y.H.C.; Project Administration, K.C., J.W.G., Y.H.C.; Funding Acquisition, E.A.B., Y.H.C., and J.W.G.

---

**Supplementary Figure 1:**
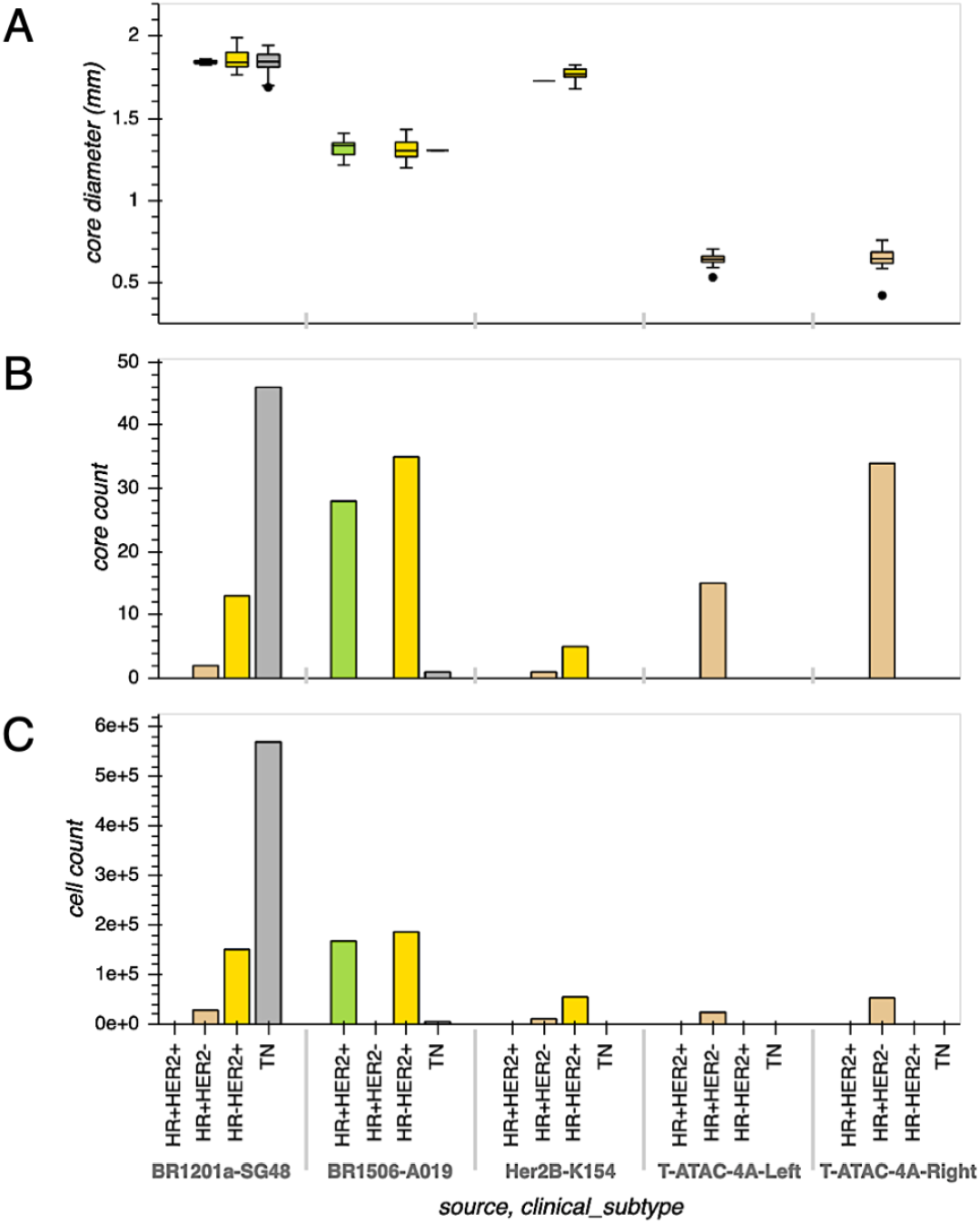
Overview of TMA composition and CyCIF panel. (A) Core diameters split by TMA source and BC subtype. (B) Core count split by TMA source and BC subtype. (C) Cell count split by TMA source and BC subtype.

**Supplementary Figure 2:**
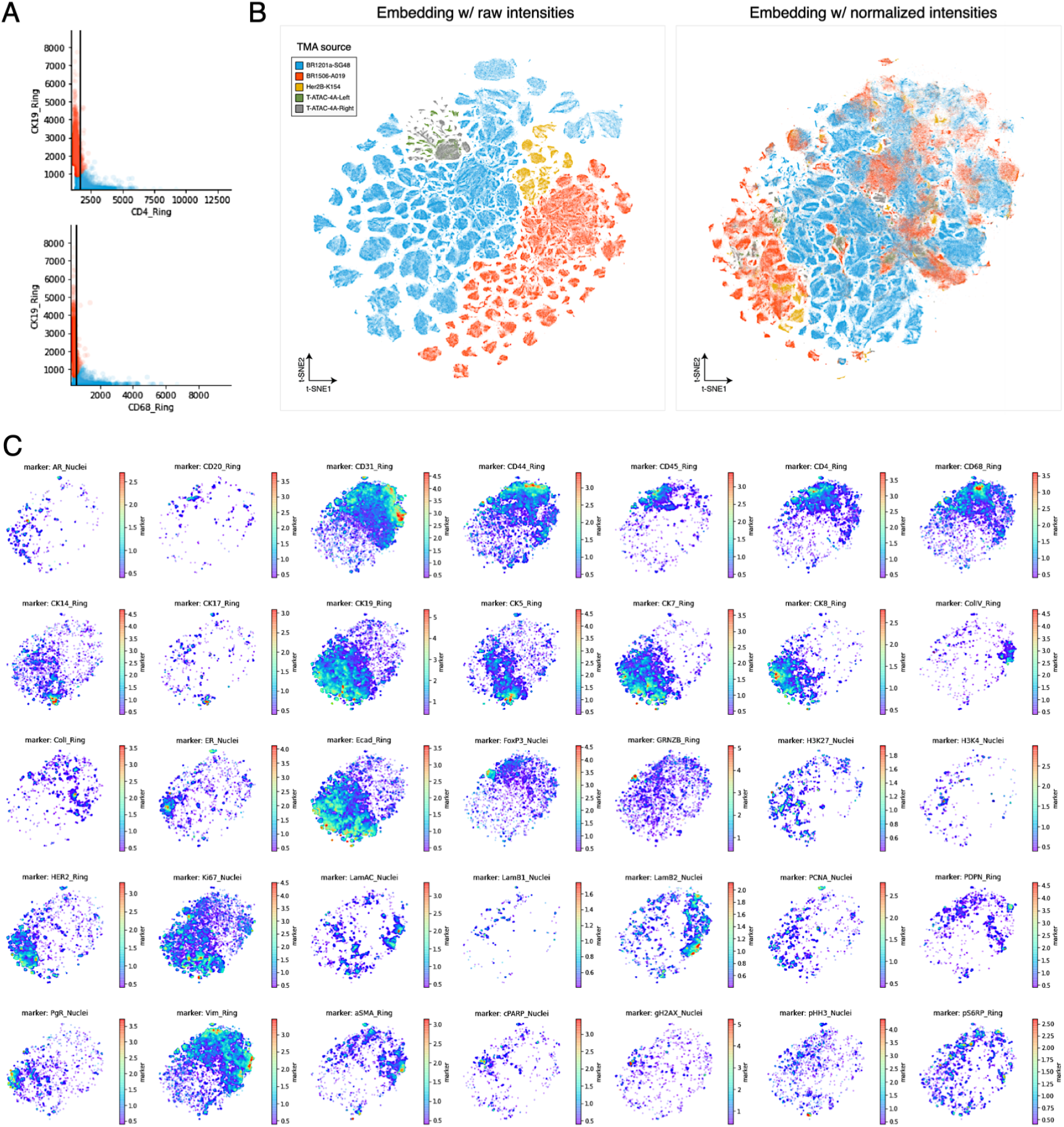
Cell mean intensity normalization across TMAs. (A) Example of RESTORE normalization [7] of CD4 and CD68 cell mean intensities for a single TMA core. Cell mean intensities of these immune markers are plotted against the cell mean intensity of epithelial CK19, a mutually exclusive marker. Cells are partitioned into positive (blue) and negative (red) populations. Black lines represent the computed normalization factors. Cells to the right of each line are above the background intensity level for that immune marker. (B) *t*-SNE embeddings of all cells using either raw (left) or normalized (right) cell mean intensities for all markers. Cells are colored according to the TMA from which they originate. A strong batch effect is observed before normalization, leading to partitioning according to TMA of origin. Following normalization, cell phenotypes shared between TMAs are co-localized. However, some TMA-specific partitioning remains due to subtype-specific marker bias within TMAs. (C) *t*-SNE normalized embedding, faceted by marker, showing only cells that have a mean intensity above the normalization factor for that marker. Color scale is log-transformed normalized intensity.

**Supplementary Figure 3:**
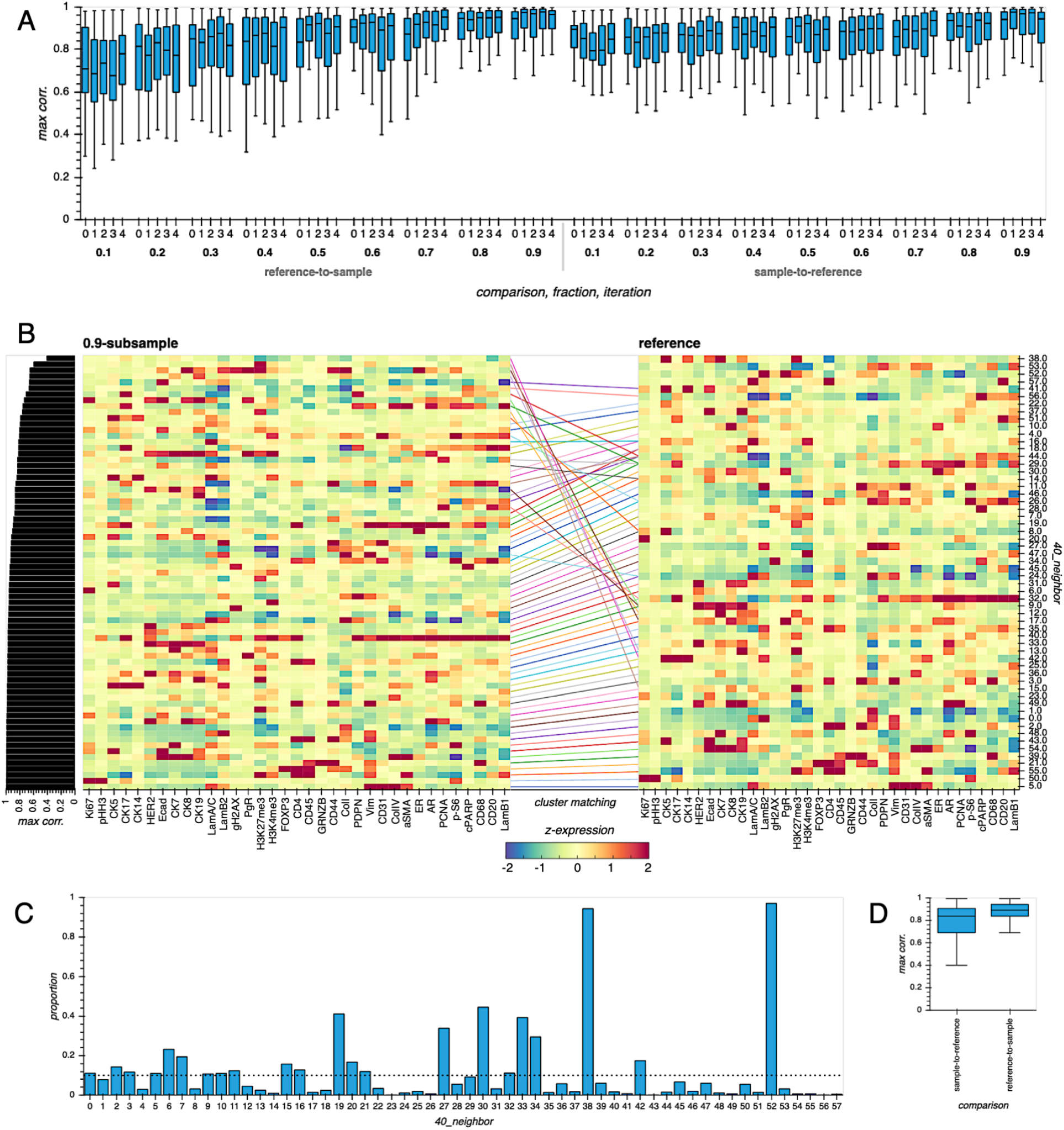
Validation of BC cell phenotype robustness. (A) Maximum Pearson’s correlations between full reference cell phenotypes and those derived using PhenoGraph on cells from a random fractional sample of TMA cores, iterated five times at each fraction level. (B) Phenotype matching between the full reference phenotypes and those derived from a 90%-subsampled fraction of TMA cores. Phenotypes are ordered by increasing matching correlation. Matching phenotypes are linked by a line, and lines are colored to discriminate between adjacent or overlapping links. 40_neighbor represents the PhenoGraph cluster labels since we set *k* = 40 when defining the *k*-nearest neighbor graph in the PhenoGraph routine. The colorbar indicates *z*-scored marker expression. (C) The proportion of cells from each reference phenotype that correspond to the 10% of TMA cores held out from the 90%-subsampled fraction from (B). Unmatched phenotypes in the full reference correspond in part to cells from held-out cores. The dotted line marks the 10% threshold. (D) Maximum Pearson’s correlation between full reference phenotypes and those derived using PhenoGraph on normalized mean intensities from all TMA cores, but with ±20% random noise added to each marker in each core.

**Supplementary Figure 4:**
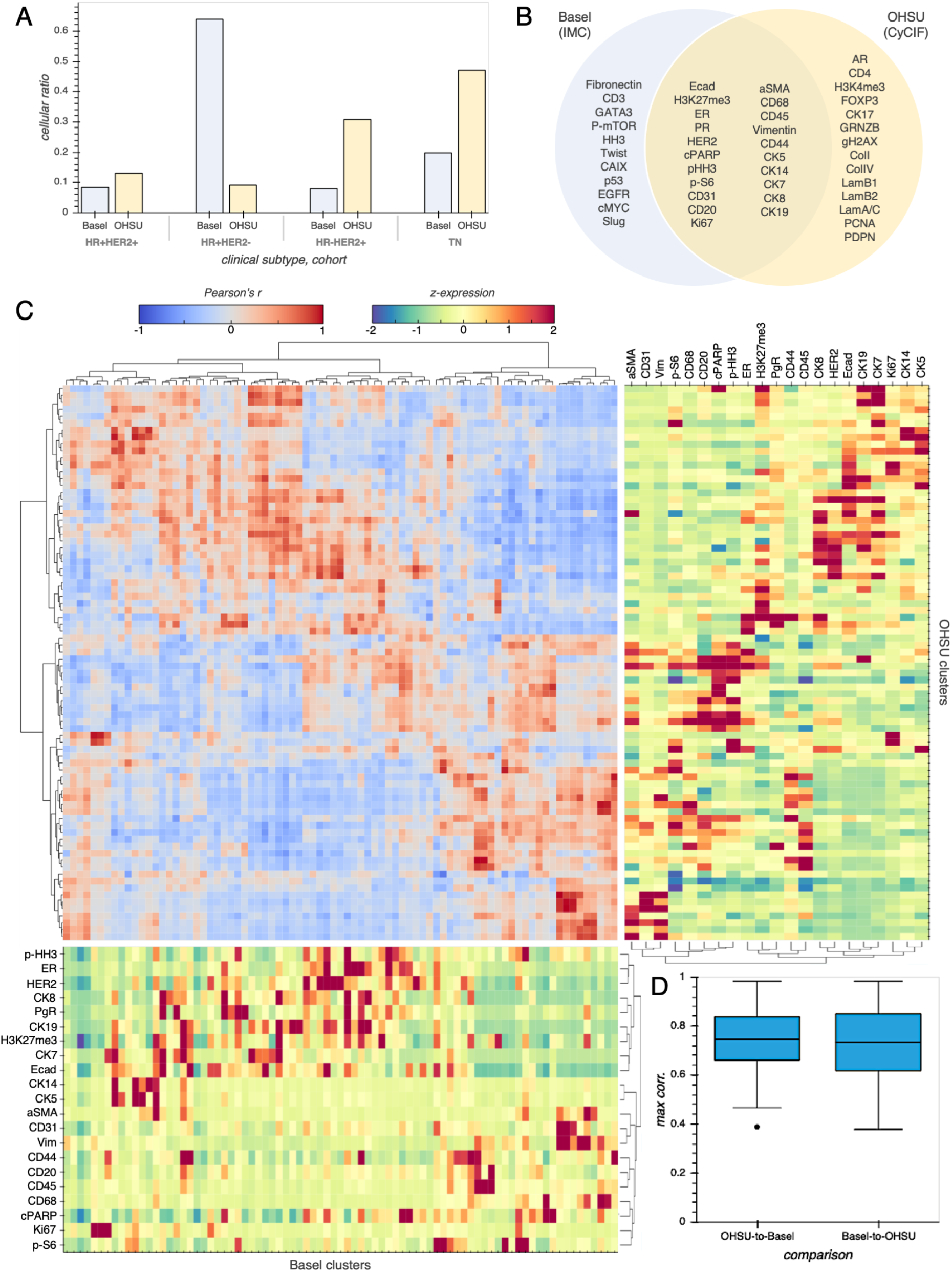
Cross-platform benchmarking of BC cell phenotypes. (A) Cellular ratio highlighting compositional differences between Basel [12] and OHSU (this work) cohorts with respect to BC subtype. (B) The intersection of the IMC and CyCIF marker panels used to stain tissues from the Basel and OHSU cohorts, respectively. (C) PhenoGraph cluster matching between Basel and OHSU cohorts. Using only the intersecting markers, cells from each cohort were independently clustered using PhenoGraph with the same parameterization, then cohort clusters were pairwise correlated and hierarchically clustered based on the resulting correlation structure. We identified highly-correlated clusters between cohorts, including those corresponding to epithelial, immune, stromal, endothelial, and proliferating cell populations. (D) Maximum Pearson’s correlation corresponding to inter-cohort cluster matches.

**Supplementary Figure 5:**
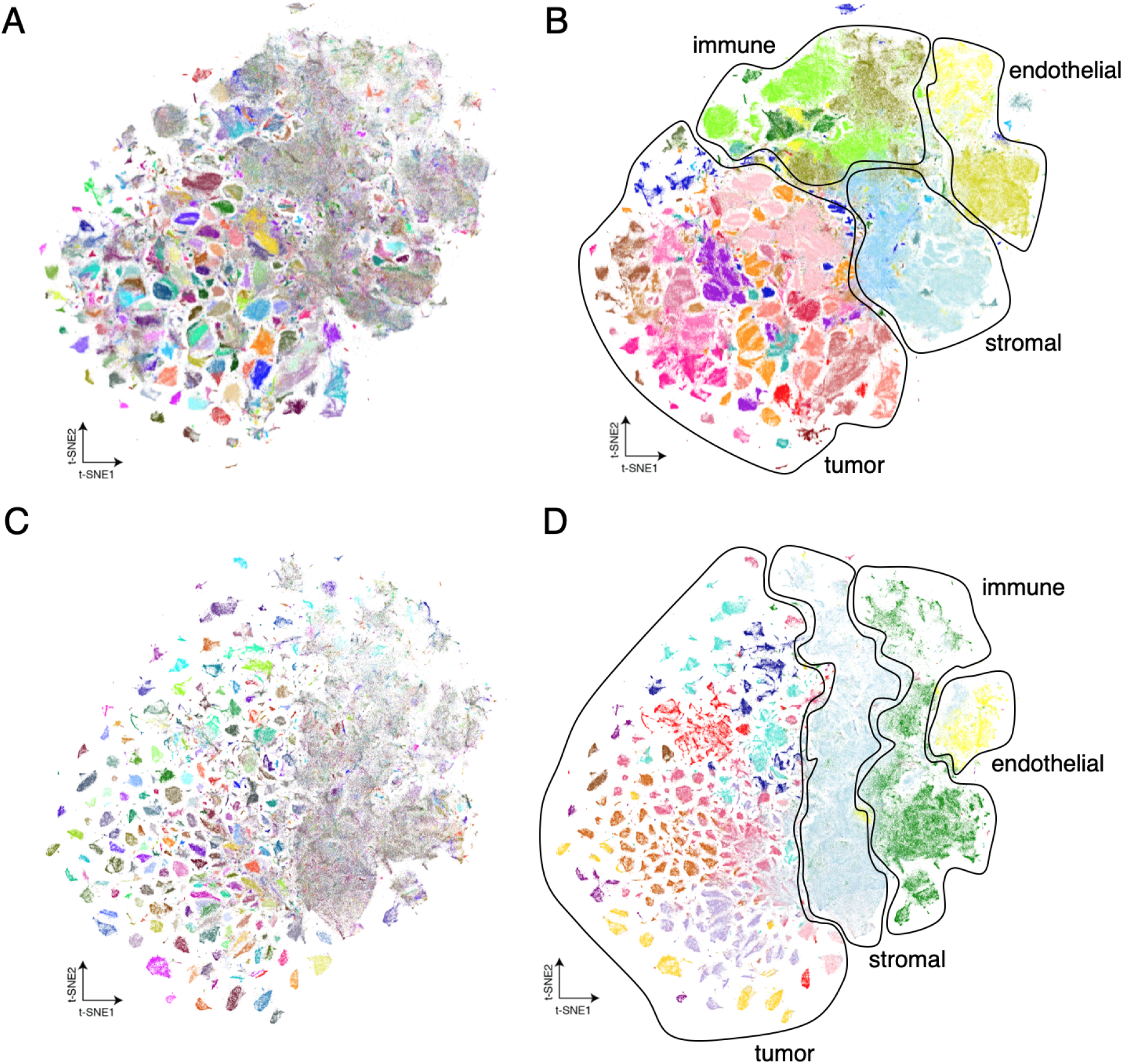
For both OHSU and Basel cohorts, tumor cells differ more between samples than immune, stromal, or endothelial cells. (A) The same *t*-SNE embedding of the OHSU dataset from Figure 1B, but with cells colored based on the unique tissue core from which they are derived. (B) The same *t*-SNE embedding of the OHSU dataset from Figure 1B, but with annotations indicating immune, stromal, endothelial, and tumor phenotypic regions. (C) A *t*-SNE embedding of the Basel dataset [12] derived using the same parameters as were used for the OHSU *t*-SNE embedding, with cells colored based on the unique tissue core from which they are derived. (D) The same *t*-SNE embedding as in (C), but with annotations indicating immune, stromal, endothelial, and tumor phenotypic regions.

**Supplementary Table 1:** Antibody panel used for CyCIF staining of tissues.

| Marker | Description | Vendor | Vendor # | Dye | Clone |
| --- | --- | --- | --- | --- | --- |
| CD20 | B cell | Abcam | ab198941 | AF488 | EP459Y |
| CD4 | T cell | Abcam | ab196147 | AF647 | EPR6855 |
| CD44 | cell adhesion | Abcam | ab216647 | AF750 | EPR1013Y |
| CD45 | lymphocyte | Abcam | ab214437 | AF750 | EP322Y |
| CD68 | macrophage | Biolegend | 916104 | AF555 | KP1 |
| FOXP3 | regulatory T cell | Biolegend | 320102 | AF750 | 206D |
| GRNZB | proteolysis | Abcam | ab219803 | AF750 | EPR20129-217 |
| CK5 | basal cytokeratin | Abcam | ab193894 | AF488 | EP1601Y |
| CK14 | basal cytokeratin | Abcam | ab212547 | AF555 | LL002 |
| CK17 | basal cytokeratin | Abcam | ab185032 | AF488 | EP1623 |
| CK7 | luminal cytokeratin | Abcam | ab185048 | AF488 | EPR1619Y |
| CK8 | luminal cytokeratin | Abcam | ab192467 | AF488 | EP1628Y |
| CK19 | luminal cytokeratin | Biolegend | 628502 | AF750 | A53-B/A2 |
| Ecad | cell adhesion | Abcam | ab201499 | AF750 | EP700Y |
| AR | hormone receptor | Sigma | 06-680-AF555 | AF555 | polyclonal |
| ER | hormone receptor | Abcam | ab205851 | AF647 | EPR4097 |
| PgR | hormone receptor | Abcam | ab199455 | AF750 | YR85 |
| HER2 | receptor tyrosine kinase | Santa Cruz | sc-33684 | AF555 | 3B5 |
| aSMA | myoepithelia | Santa Cruz | sc-32251 | AF488 | 1A4 |
| CD31 | endothelia | Abcam | ab218582 | AF647 | EPR3094 |
| Vimentin (Vim) | mesenchyme | CST | 9854 | AF488 | D21H3 |
| ColI | extracellular matrix | Abcam | ab215969 | AF750 | EPR7785 |
| ColIV | extracellular matrix | ThermoFisher/eBioscience | 51-9871-82 | AF647 | 1042 |
| LamA/C | nuclear structure | Sigma | SAB4200236 | AF750 | 4C11 |
| LamB1 | nuclear structure | Abcam | ab194106 | AF488 | EPR8985(B) |
| LamB2 | nuclear structure | Abcam | ab200427 | AF647 | EPR9701(B) |
| H3K4me3 | epigenetic activation | CST | 11960 | AF555 | C42D8 |
| H3K27me3 | epigenetic repression | CST | 5499 | AF488 | C36B11 |
| PDPN | migration signalling | Biolegend | 916606 | AF555 | polyclonal |
| cPARP | apoptosis | CST | 6894 | AF555 | D64E10 |
| gH2AX | replication stress | Abcam | ab195189 | AF647 | EP854(2)Y |
| Ki67 | proliferation | CST | 12075 | AF647 | D3B5 |
| PCNA | proliferation | CST | 8580 | AF488 | PC10 |
| pHH3 | mitosis | CST | 3465 | AF488 | D2C8 |
| p-S6 | translational activation | CST | 3985 | AF555 | D57.2.2E |

**Supplementary Table 2:**
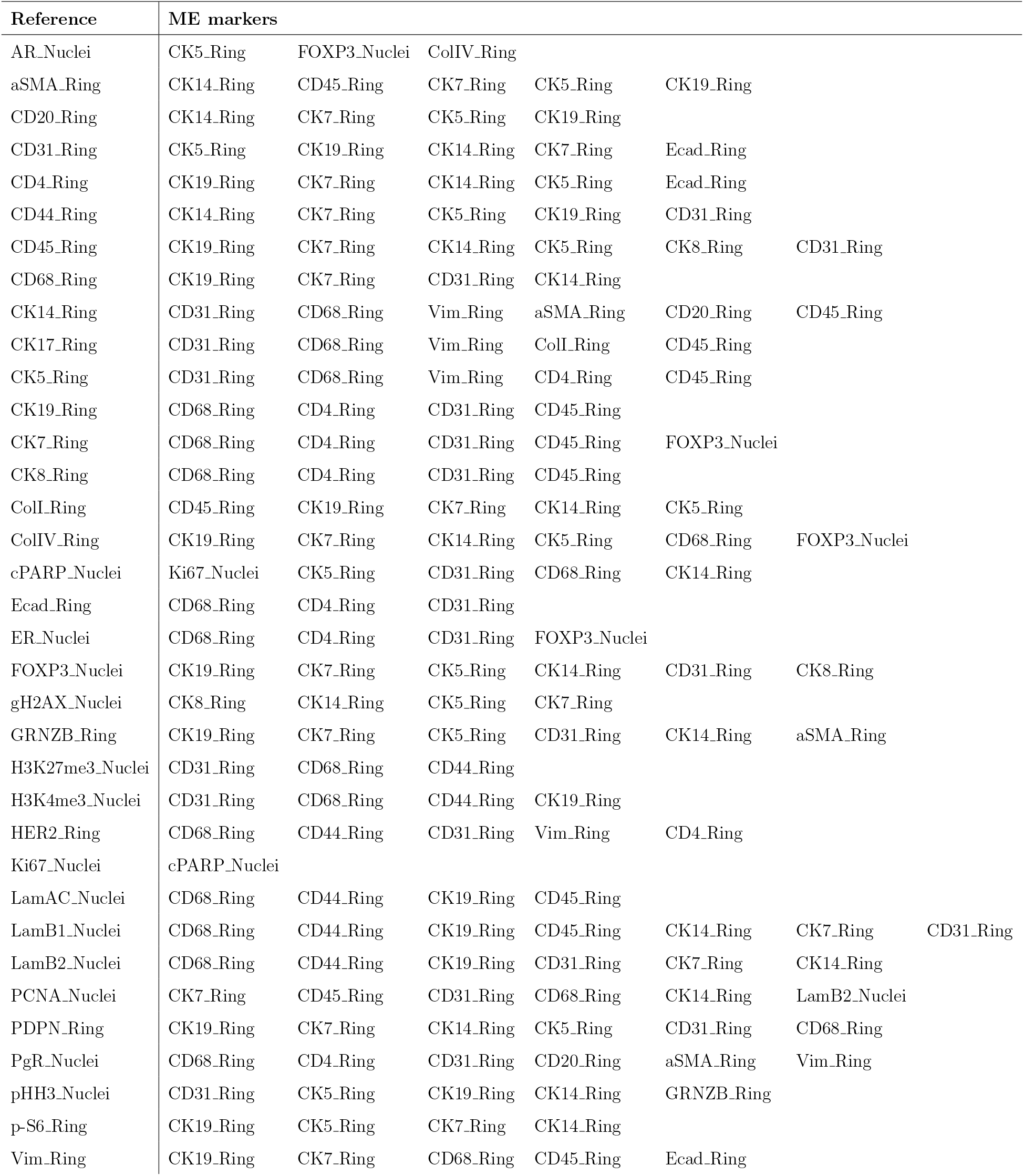
Putative reference and mutually-exclusive (ME) marker pairs used for RESTORE normalization of cell mean intensities. Each marker name indicates from which compartment its mean intensity was extracted. “Ring” indicates that a marker’s intensity was extracted from the ring-shaped cytoplasmic segmentation masks derived by subtracting the “Nuclei” segmentation masks from “Cell” segmentation masks.

